# RUNX/CBFβ transcription factor complexes promote the phenotypic plasticity of metastatic breast cancer cells

**DOI:** 10.1101/562538

**Authors:** Ran Ran, Hannah Harrison, Nur Syamimi Ariffin, Rahna Ayub, Henry J Pegg, Wensheng Deng, Andrea Mastro, Penny D. Ottewell, Anna M. Fowles, Susan M. Mason Karen Blyth, Ingunn Holen, Paul Shore

## Abstract

Epithelial to mesenchymal transition (EMT) is a dynamic process that drives cancer cell plasticity and is thought to play a major role in metastasis. Here we show that the plasticity of metastatic breast cancer cells can be promoted by the activity of the RUNX transcription factors. We demonstrate that the RUNX co-regulator CBFβ is essential to maintain the mesenchymal phenotype of triple-negative breast cancer cells and that CBFβ-depleted cells undergo a mesenchymal to epithelial transition (MET) and re-organise into acini-like structures, reminiscent of those formed by epithelial breast cells. We subsequently show, using an inducible CBFβ system, that the MET can be reversed, thus demonstrating the plasticity of RUNX/CBFβ-mediated EMT. Moreover, the MET can be reversed by expression of the EMT transcription factor Slug whose expression is dependent on CBFβ, RUNX1 and RUNX2. Finally, we demonstrate that loss of CBFβ inhibits the ability of metastatic breast cancer cells to invade bone cell cultures and suppresses their ability to form bone metastases *in vivo*. Together our findings demonstrate that the RUNX/CBFβ complexes can determine the plasticity of the metastatic cancer cell phenotypes, suggesting that their regulation in different micro-environments may play a key role in the establishment of metastatic tumours.

## The role of CBFβ in Breast Cancer

The triple-negative sub-type of breast cancer is a highly aggressive cancer for which treatment options are limited^1^. Expression of the RUNX transcription factors in patients with the triple-negative subtype of breast cancer correlates with a poor prognosis^2 3^. Emerging evidence suggests that the role of RUNX proteins in breast cancer is dependent on the specific RUNX factor involved and the sub-type of breast cancer cell^4^. In order to consider the RUNX factors as viable targets in breast cancer therapies it is therefore critically important to determine the role of the different factors in different sub-types of breast cancer. Since CBFβ facilitates the function of all three RUNX transcription factors, establishing its role in determining the phenotype of breast cancer cells is essential^5 6^.

Mutations in CBFβ are amongst the most frequently reported for breast cancer tumours, suggesting a tumour suppressor role for CBFβ in ER+ breast cancer^7 8^. In contrast, we and others have previously shown that expression of RUNX2 and CBFβ contribute to the metastatic phenotype of triple negative breast cancer cells^9 10 11 12^. In this context it is therefore the maintained expression of RUNX factor activity that promotes their metastatic phenotype.

Epithelial to mesenchymal transition (EMT) contributes to the progression of metastatic cancer as it enables cancer cells to become migratory and invasive^13 14 15^. EMT is also a plastic program in which cells dynamically transition along a continuum of states between epithelial and mesenchymal phenotype^16 17^. This plasticity is thought to enable mesenchymal cancer cells to switch from a migratory to an epithelial phenotype during colonization of the metastatic niche. Previous studies have shown that RUNX factors have different roles in EMT in different breast cell-types^18 19 20 21^. In ER-negative MCF10A cells, expression of RUNX1 suppresses EMT, whereas RUNX2 stimulates an EMT-like phenotype in these cells. In triple negative MDA-MB-231 cells, depletion of RUNX2 inhibited their invasive capacity, consistent with suppression of EMT^22^.

Since CBFβ forms functional complexes with all RUNX transcription factors, it is essential to establish its role in breast cancer metastasis. Here we show that CBFβ-depleted triple negative cells were able to undergo a reversion from a mesenchymal phenotype to an epithelial phenotype. Indeed, acini-like structures with clear lumen were observed, reminiscent of those formed by normal epithelial breast cells. Remarkably, re-induction of CBFβ, using an inducible system, led to a complete reversion from the epithelial phenotype to a mesenchymal phenotype, demonstrating the plasticity of CBFβ-driven EMT. We subsequently show that CBFβ maintains the mesenchymal phenotype through activation of the EMT transcription factor Slug. We also show that the maintenance of the mesenchymal phenotype is dependent upon CBFβ, RUNX1 and RUNX2, as depletion of any of these factors resulted in MET. Finally, we demonstrate that loss of CBFβ, inhibits the ability of metastatic breast cancer cells to develop bone metastases *in vivo*. Together our findings demonstrate that the RUNX/CBFβ complexes can determine the plasticity of the metastatic cancer cell phenotypes, suggesting that their regulation in different micro-environments may play a key role in the establishment of metastatic tumours.

## Results

### CBFβ maintains the mesenchymal phenotype of metastatic breast cancer cells

We have previously shown that depletion of CBFβ in metastatic MDA-MB-231 breast cancer cells inhibits their ability to migrate^11^. This is not restricted to MDA-MB-231 cells as knockdown of CBFβ in the MDA-MB-468 metastatic cell line also inhibited their migration (Fig. S1). We also observed that CBFβ-depleted cells exhibit a more rounded phenotype and were unable to migrate in a wound-healing assay (Fig. 1A). These observations suggested that the cells had undergone a profound phenotypic change. When grown in 3D culture, normal mammary epithelial cells differentiate into organized acini structures with clearly identifiable lumen, reminiscent of normal mammary gland morphology, whereas metastatic cells exhibit a mesenchymal phenotype and form a stellate appearance. We therefore compared the 3D morphology of wild-type metastatic breast cancer cells, MDA-MB-231, with MDA-MB-231 cells depleted of CBFβ. As expected, when grown in 3D, MDA-MB-231 cells spread throughout the culture, displaying a typical stellate pattern that reflects their mesenchymal phenotype (Fig.1B) ^23^. In contrast, depletion of CBFβ resulted in a striking loss of the stellate pattern and the formation of clusters and spherical colonies (Fig. 1B).

**Figure 1:**
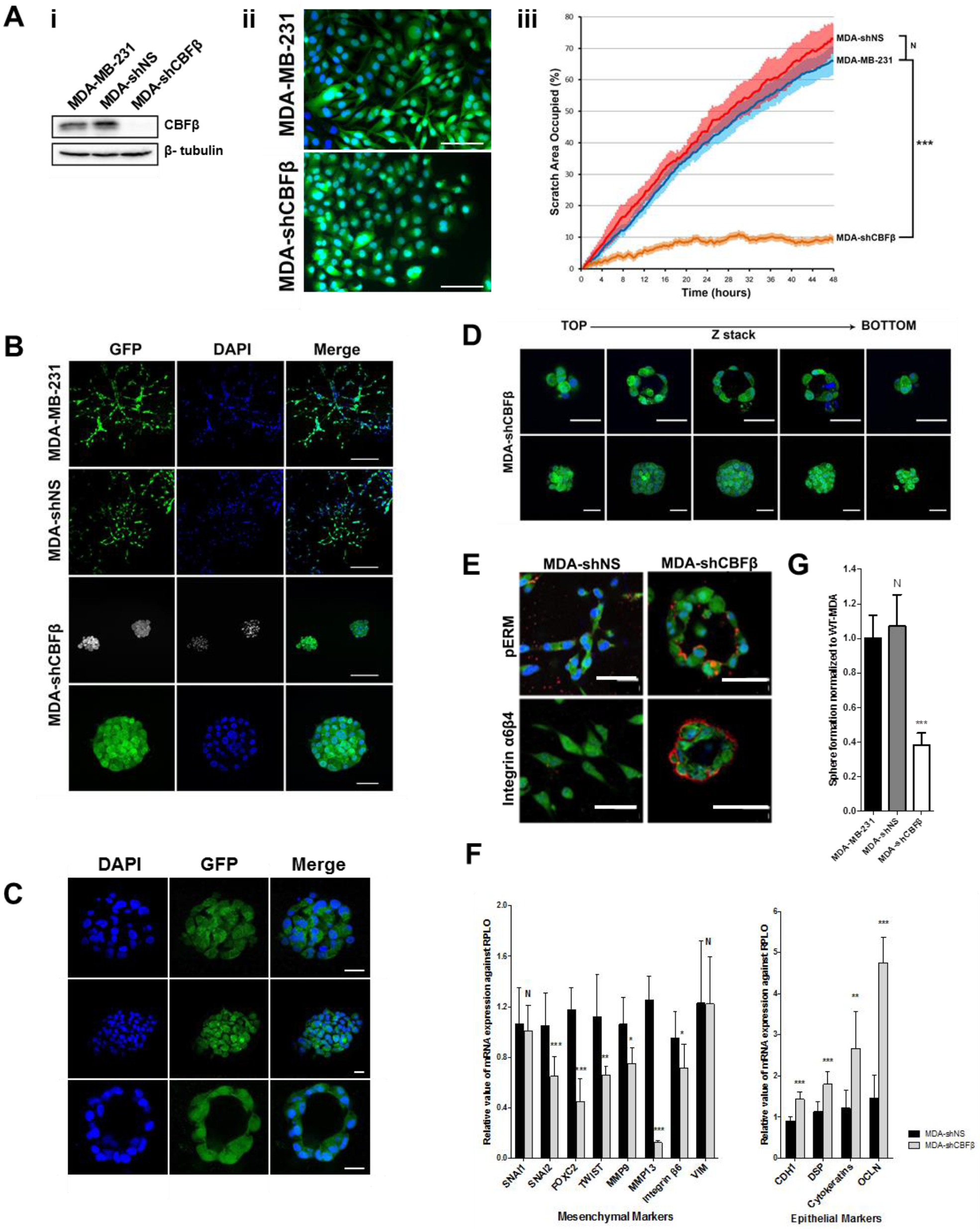
CBFβ maintains the mesenchymal phenotype of metastatic breast cancer cells. (A) CBFβ loss results in phenotypic changes. i) Western blot shows knockdown of CBFβ in MDA-MB-231 cells, MDA-shNS and MDA-shCBFβ. β-Tubulin was used as a loading control. ii) Fluorescent microscopy comparing the shape of GFP-expressing MDA-MB-231 cells and CBFβ-depleted cells (MDA-shCBFβ). iii) Following scratch to monolayer, live images were taken every 20 mins for 48 hours. Graph shows significant reduction in cell mobility in the absence of CBFβ. Statistical analysis performed using ANOVA. (B) Loss of CBFβ results in cluster formation in 3D culture. MDA-MB-231, MDA-shNS and MDA-shCBFβ were grown in 3D Matrigel for 14 days. GFP (green) was stably expressed in all cell lines. Cells were fixed and nuclei stained with DAPI (blue). The wild-type and MDA-shNS cells showed a stellate mesenchymal growth pattern whilst MDA-shCBFβ cells formed discrete clusters. Scale bars are 200μm in top three rows and 50μm in row four. (C) Confocal microscopy showing different sizes of clusters in the MDA-shCBFβ cells. The lower panels clearly show a smaller cluster that resembles an acinus. Scale bars are 25μm. (D) Z-stack analysis showing that some smaller clusters contain lumen and some do not. Scale bars are 50μm. (E) Acini express polarity markers. Immunofluorescence microscopy of MDA-shNS and MDA-shCBFβ cells in 3D Matrigel after 14 days. Cells were fixed and stained for apical or basement markers including phospho Ezrin/Radixin/Moesin (pERM) and Integrins (Red) and nuclei were stained with DAPI (Blue). Scale bars are 50μm. (F) Loss of CBFβ causes a reduction in mesenchymal markers and increased expression of epithelial markers. qRT-PCR was performed for epithelial and mesenchymal markers in MDA-shNS or MDA-shCBFβ. (G) Loss of CBFβ causes a reduction in the number of mammosphere-forming colonies. Mammospheres were counted after 5 days growth in non-adherent culture.

We next used confocal microscopy to determine the extent to which the cells had differentiated in the absence of CBFβ (Fig. 1C & 1D). Remarkably, a small number of these structures contained clearly identifiable lumen reminiscent of acini typically formed by mammary epithelial cells (Fig 1. C-E), whilst the larger clusters did not appear to contain lumen (Fig. 1D, lower panels). Subsequent immunofluorescence staining demonstrated that some of these structures were also polarised and expressed the basement membrane markers integrins α6 and β4 on the apical surface and phospho-ezrin-radixin-moesin (pERM) on the luminal surface, as found in wild-type acini structures (Fig. 1E)^24^. These observations suggested that the cells had actually undergone a mesenchymal to epithelial transition (MET). To confirm this, we used RT-PCR to determine the expression of a number of mesenchyme and epithelial markers in wild-type MDA-MB-231 cells and the MDA-shCBFβ cells. Of the eight mesenchymal markers analysed, seven had significantly reduced expression in the MDA-shCBFβ cells. This reduction in mesenchymal markers was accompanied by an increase in expression of the four epithelial markers analysed (Fig. 1F).

Another feature of EMT is that it leads to an increase in the number of cancer stem cells (CSCs)^17^. The potential for stem cell formation can be monitored using the mammosphere assay^25^. We therefore examined the number of mammospheres formed in the presence and absence of CBFβ. Loss of CBFβ resulted in a significant reduction in the number of mammospheres formed when compared to parental MDA-MB-231 cells, suggesting that less in the absence of CBFβ the potential to produce CSCs is reduced (Fig. 1G).

Taken together, our data demonstrate that CBFβ is essential for the maintenance of the mesenchymal phenotype in MDA-MB-231 cells and suggests that loss of CBFβ expression causes differentiation towards a more epithelial-like phenotype.

### CBFβ regulates EMT/MET state-transition

Dynamic EMT/MET state-transition is thought to enable metastatic cancer cells to switch between the two phenotypes, a property that allows them to both invade and colonise other tissues^17 26^. We therefore sought to establish if the dynamic EMT/MET state-transition could be regulated by the expression of CBFβ. To do this we created a cell line in which the activity of CBFβ can be regulated by the addition of 4OH-T (Fig. 2Ai)^27^. MDA-shCBFβ cells stably expressing a CBFβ-ER fusion protein were generated and western blotting confirmed that the addition of 4OH-T to these cells induced nuclear expression of CBFβ-ER (Fig. 2Aii). RT-PCR analysis showed that there was also a concomitant increase in expression of the known RUNX-target genes OPN, MMP9 and MMP13 (Fig. 2B). Activation of CBFβ also rescued the migration defect, as determined by a significant increase in the capacity of the cells to invade Matrigel and migrate in a wound-healing assay in the presence of 4OH-T (Fig. 2C). Microscopic analysis of the cells in 3D culture, in the presence and absence of 4OH-T, revealed a remarkable difference in phenotypes. In the absence of 4OH-T the cells formed clusters as expected (Fig. 2D, upper panel). However, cells grown in the presence of 4OH-T had spread throughout the culture and were indistinguishable from the parental MDA-MB-231 cells (Fig. 2D, see Fig. S2 for controls). Furthermore, activation of CBFβ induced an increase in expression of mesenchymal markers and a decrease in epithelial markers (Fig. 2E). Consistent with this, we observed an increase in the number of mammospheres formed following culture with 4OH-T (Fig. 2F). To demonstrate that CBFβ can dynamically regulate the plasticity of the cells we added 4OH-T to MDA-CBFβ-ER cells that had already formed clusters in 3D culture (Fig. 2G). In the absence of 4OH-T the clusters continued to grow with little dispersion. In contrast, after three days in the presence of 4OH-T branched structures were observed, and after six days cells were clearly dispersing (Fig. 2G). These data show that CBFβ drives the mesenchymal phenotype of MDA-MB-231 and that the EMT/MET state-transition can be dynamically regulated in metastatic cells by CBFβ.

**Figure 2:**
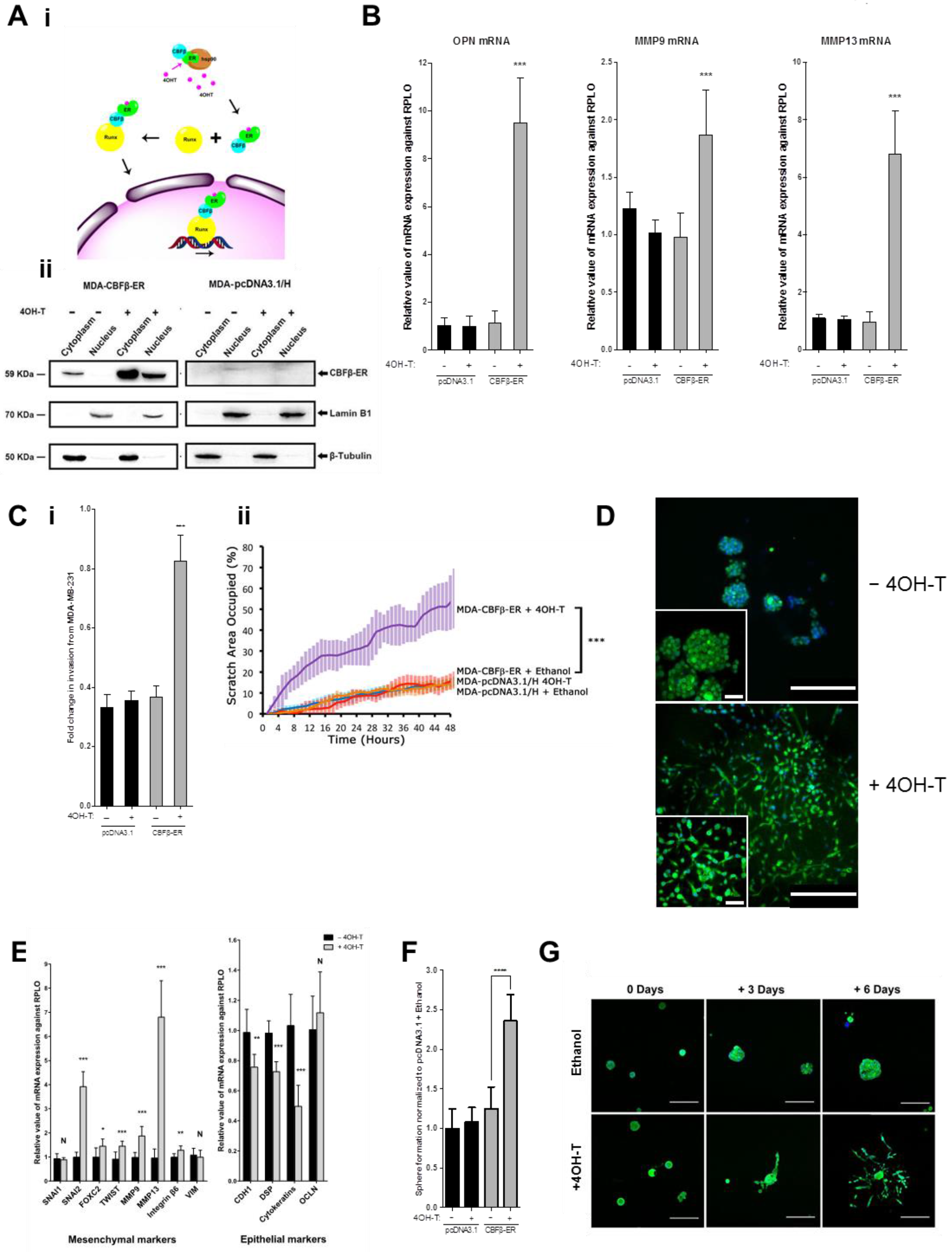
CBFβ regulates EMT/MET state-transition. (A) Inducible activation of CBFβ. i) Schematic showing inducible cell line in which a CBFβ-ER fusion protein, encoding mouse CBFβ, was stably expressed in MDA-shCBFβ cells. Incubation with 4OH-T causes CBFβ-ER to translocate to the nucleus. ii) Western blot showing nuclear and cytoplasmic extracts from 4OH-T and ethanol treated MDA-CBFβ-ER cells using anti-CBFβ antibody. Lamin B1 and Tubulin were used as nuclear and cytoplasmic loading controls respectively. (B) RUNX target genes are activated in MDA-CBFβ-ER cell lines by 4OH-T. Total RNAs were used to analyse the mRNA levels of known RUNX target genes, OPN (osteopontin), MMP9 (matrix metallopeptidase-9) and MMP-13 (matrix metallopeptidase-13). (C) Activation of CBFβ-ER with 4OH-T restores cell invasion and mobility capacity. i) Matrigel invasion assay showing the recovery of invasion in the presence of 4OH-T following 24 hours culture. ii) Following scratch to monolayer, live images were taken every 20 mins for 48 hours. Graph shows migration capacity is restored in the presence of 4OH-T in CBFβ-ER cells. Statistical analysis performed using ANOVA. (D) Induction of CBFβ-ER with 4OH-T restores the mesenchymal phenotype in 3D culture. The GFP expressing cells were visualised by fluorescence microscopy and DAPI staining. The upper panel shows MDA-CBFβ-ER cells grown after addition of vehicle (ethanol). The lower panel shows 4OH-T treated MDA-CBFβ-ER cells. Scale bars are 200μm in large image and 50μm in inset image. (E) Re-expression of CBFβ restores EMT marker gene expression. Epithelial and mesenchymal marker genes were analysed by qRT-PCR on RNA isolated from MDA-CBFβ-ER cells grown in the presence 4OH-T or ethanol. (F) Induction of CBFβ-ER restores mammosphere formation capacity. MDA-CBFβ-ER and control MDA-pcDNA3.1/H cells were grown with 4OH-T or ethanol and their mammopshere forming capacity subsequently determined. Mammospheres were counted after 5 days growth in non-adherent culture. (G) Immunofluorescence microscopy of MDA-CBFβ-ER cells in 3D culture showing dispersion of cell clusters upon activation of CBFβ. Cells were grown in Matrigel for 8 days (day 0 of 4OH-T addition), prior to addition of 4OH-T or ethanol. Cells were fixed and visualised for GFP (green) and DAPI (blue). Scale bars are 200μm.

### Depletion of RUNX1 and RUNX2 induces MET

Since both RUNX1 and RUNX2 are expressed in MDA-MB-231 cells we sought to establish if a double knockdown of RUNX1 and RUNX2 phenocopied the effect of CBFβ-knockdown. We therefore generated cell lines in which either RUNX1 or RUNX2 were depleted and a cell line in which both proteins were depleted (Fig. 3A; S3A & S4A). All three cell lines displayed a significant decrease in migration and invasion capacity (Figs. 3B & C; S3 & S4) as well as a decrease in mammosphere formation (Figs. 3D; S3 & S4). Mesenchymal and epithelial marker expression confirmed that each of the knockdown cell lines had undergone MET (Figs. 3E; S3 & S4). All of the lines formed clusters in 3D culture (Figs. 3F; S3 & S4). Additionally, we observed acini-like structures in the double knockdown cells demonstrating that depletion of both proteins does indeed phenocopy the effect of CBFβ loss (Fig. 3F). We did not observe acini in the single knockdown cell lines, although acini-like structures have previously been reported in RUNX2-knockdown cells^22^. Taken together, these findings demonstrate that CBFβ, RUNX1 and RUNX2 are necessary for the maintenance of the mesenchymal phenotype in MDA-MB-231 cells.

**Figure 3.**
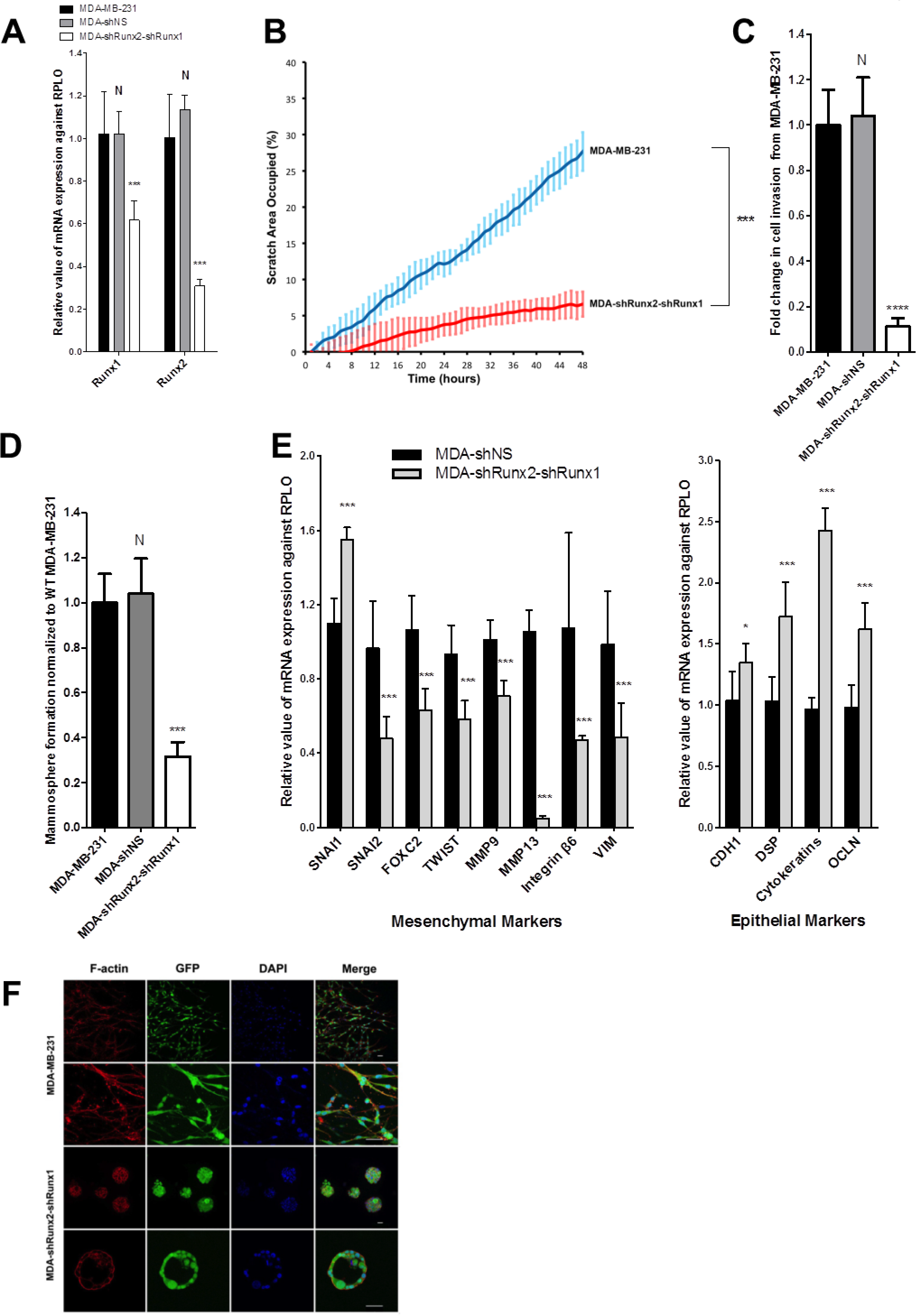
Knockdown of RUNX1 and RUNX2 induces MET. (A) qRT-PCR showing knockdown of RUNX1 and RUNX2 by shRNA. Total RNAs were used to analyse the mRNA levels of RUNX1 and RUNX2. RPLO mRNA was used as a control for normalisation and relative values are shown. (B) Scratch assays showing inhibition of migration in MDA-shRUNX2-shRUNX1 cells. Following scratch to monolayer, live images were taken every 20 mins for 48 hours. Graph shows significant reduction in cell mobility in the absence of RUNX1/RUNX2. (C) Matrigel invasion assay showing inhibition of migration in MDA-shRUNX2-shRUNX1 cells. (D) Loss of RUNX1 and RUNX2 causes a reduction in mammosphere formation. Mammospheres were counted after 5 days growth in non-adherent culture. (E) Loss of RUNX1 and RUNX2 in MDA-MB-231 cells causes a reduction in mesenchymal markers and increased expression of epithelial markers as seen by RT-PCR. (F) Loss of RUNX1 and RUNX2 results in cluster formation in 3D culture. Immunofluorescence microscopy of MDA-MB-231 and MDA-shRUNX2-shRUNX1 cells in Matrigel after for 14 days growth. The confocal images show equatorial cross sections. Scale bars are 50μm

### RUNX/CBFβ drives Slug expression to maintain EMT

The EMT transcription factor Slug is encoded by the SNAI2 gene whose expression is reduced in CBFβ, RUNX1 and RUNX2 knockdown cell lines (Figs. 1F, S3 & S4)^18 16^. To confirm that the changes in SNAI2 mRNA expression correlated with Slug protein expression we used western blotting to determine Slug expression in the MDA-CBFβ-ER cell line in the presence and absence of 4OH-T (Fig. 4A). In the absence of 4OH-T Slug expression was significantly reduced whereas addition of 4OH-T resulted in a clear increase in expression (Fig. 4A). This suggested that the reduction in Slug expression in CBFβ-depleted cells contributes to the loss of the observed EMT. To test this, we predicted that re-expression of Slug should rescue the migration phenotype of CBFβ-depleted cells. We therefore generated a cell line stably expressing Slug in the CBFβ-depleted cells. Analysis of these cells in the wound-healing assay revealed that their ability to migrate had significantly recovered (Fig. 4B). Moreover, these cells exhibited a stellate phenotype reminiscent of the wild-type MDA-MB-231 cells, when grown in 3D culture (Fig. 4C, lower panels). Furthermore, Slug was depleted in the RUNX1/2-knockdown cell line and their migratory phenotype was rescued by the ectopic expression of Slug (Fig. 4D). ChIP analysis of the SNAI2 promoter clearly demonstrated that both RUNX1 and RUNX2 could immunoprecipitate the promoter but not a control sequence 1.5kb upstream (Fig. 4E). Binding to the promoter was abrogated in RUNX1 and RUNX2-depleted cells (Fig. 4E). Taken together, these data demonstrate that the RUNX/CBFβ complexes maintain the mesenchymal phenotype of MDA-MB-231 cells, at least in part, by regulating the expression of the EMT transcription factor Slug.

**Figure 4.**
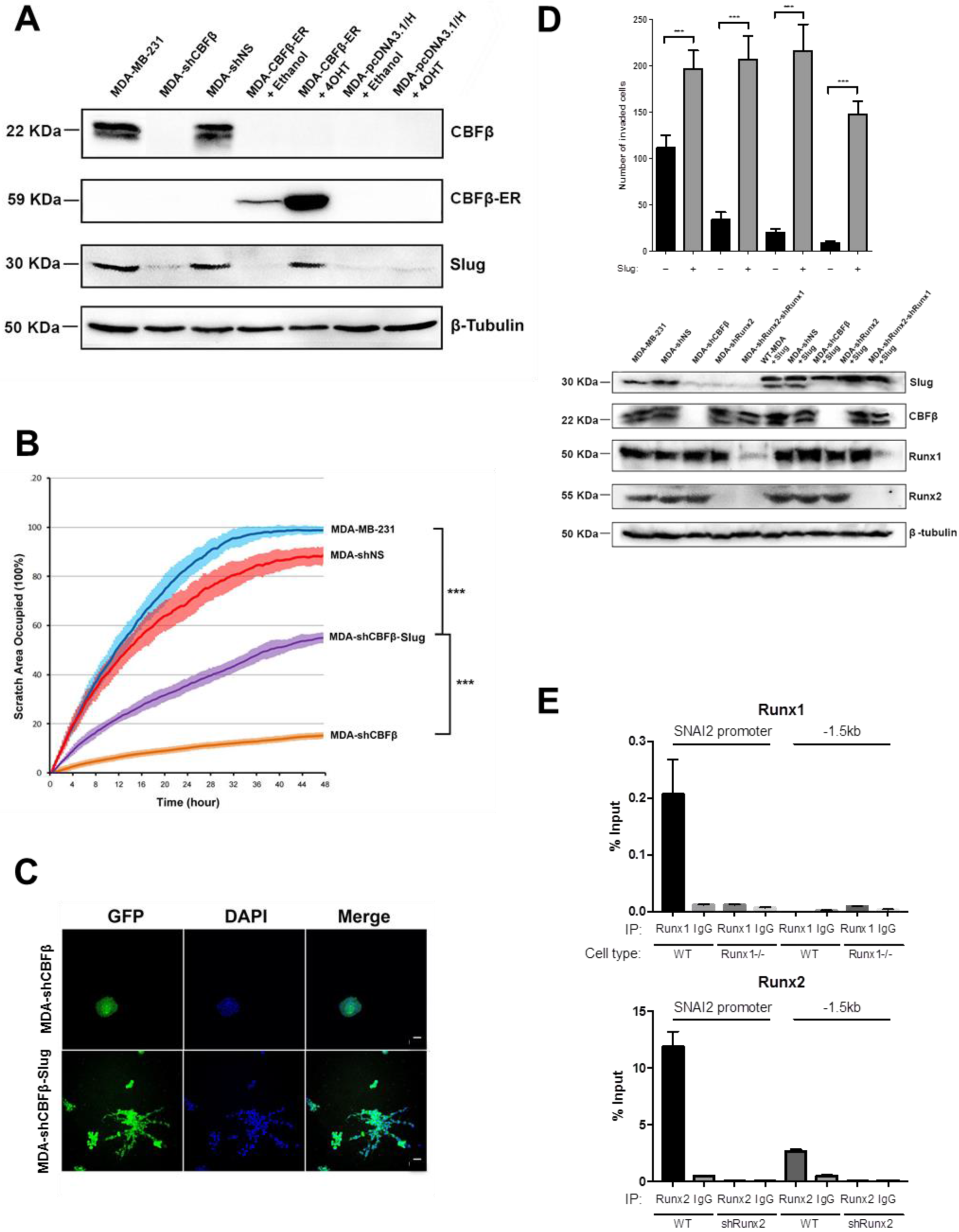
RUNX/CBFβ maintains EMT via the EMT transcription factor Slug. A) Activation of CBFβ restores expression of Slug in MDA-CBFβ-ER cells. Western blot for CBFβ and Slug performed on cell lysates from WT-MDA-MB-231, MDA-shCBFβ, MDA-shNS, MDA-CBFβ-ER and MDA-pcDNA3.1/H cells in the presence and absence of 4OH-T. Tubulin was used as loading control. (B) Slug expression in MDA-shCBFβ cells partially restores cell migration capacity. Wound healing assay showing the recovery of migration capacity in cell depleted of CBFβ and cells stably expressing Slug. Following scratch to monolayer, live images were taken every 20 mins for 48 hours. (C) Re-expression of Slug induces a mesenchymal spreading phenotype in MDA-shCBFβ cells in 3D culture. Scale bars are 50μm. (D) Expression of Slug in RUNX-depleted cells restores their invasive capacity. Cells were transfected with a Slug-Myc expressing plasmid or the control plasmid pcDNA3.1/Hygro. Western analysis shows expression of exogenous Slug-Myc. Expression of endogenous Slug (lower band in the Slug panel) and exogenous Slug-Myc (upper band in the Slug panel) was analysed by western blot from total cell lysates 72 hours after transfection. Matrigel invasion assays showing transiently transfected Slug-Myc-expressing cells. Invasion was restored in all three knockdown cells as indicated. (E) ChIP assay showing binding of RUNX1 and RUNX2 to the SNAI2 promoter. Percentage input of RUNX protein binding is shown relative to the anti-rabbit IgG antibody.

### CBFβ contributes to the development of bone metastasis in mice

To investigate the metastatic potential of the CBFβ-depleted cells we first determined their tumourigenic capacity by direct transplant into the mammary fat pad of mice (Fig. 5A)^28^. Numbers of tumours in both control and CBFβ-depleted cells were similar, however tumour volume in the CBFβ-depleted cells was significantly less than the shNS cells (Fig. 5A). This correlated with the slower growth rate of these cells observed in 3D culture (Fig. 5B). To determine if the MET induction caused by the loss of CBFβ affected the ability of cells to invade a metastatic niche we first compared the growth of parental MDA-MB-231 cells with the CBFβ-depleted cells in co-cultures with osteoblasts which had been growing for 12 weeks in 3D (Fig. 5C)^29^. Eight days after adding the cells to the osteoblast cultures, the GFP-expressing parental cells had invaded throughout the culture whereas the CBFβ-depleted cells formed discrete clusters and did not permeate the osteoblast network (Fig. 5C). Having established that the CBFβ-depleted cells retain the ability to form localised tumours but have a restricted capacity to grow in osteoblast culture we next tested their ability to establish bone metastases after injection into the circulation by intracardiac injection (Fig. 5D)^30^. Initial experiments with the CBFβ-depleted cells resulted in bone tumours in which CBFβ-expression had recovered from the shRNA (data not shown). This observation suggested that CBFβ was indeed required for bone metastases to form. We therefore created a CBFβ-depleted cell line using CRISPR-Cas9 (Figs. 5Di; S5)^31^. These CBFβ−/− cells showed the same loss of mesenchymal phenotype, decreased invasion, mammosphere formation and mesenchymal markers as seen in the shCBFβ cells (Fig S5). Intracardiac injection of the CBFβ-CRISPR cells into mice resulted in a striking reduction of the number of mice developing bone tumours. compared to control. We found that 7 out of 11 animals injected with the control cells developed tumours in the hind limbs, with numerous cancer-induced bone lesions detected, compared to 3 out of 7 animals injected with CBFβ-CRISPR (Dii). In addition, the animals receiving the control cells had higher numbers skeletal lesions (average=6.45 lesions/mouse) compared to those receiving CBFβ-CRISPR cells (average =1.28 lesions/mouse), (Fig. Di). These data demonstrate that loss of CBFβ reduces the ability of MDA-MB-231 cells to metastasise to bone.

**Figure 5.**
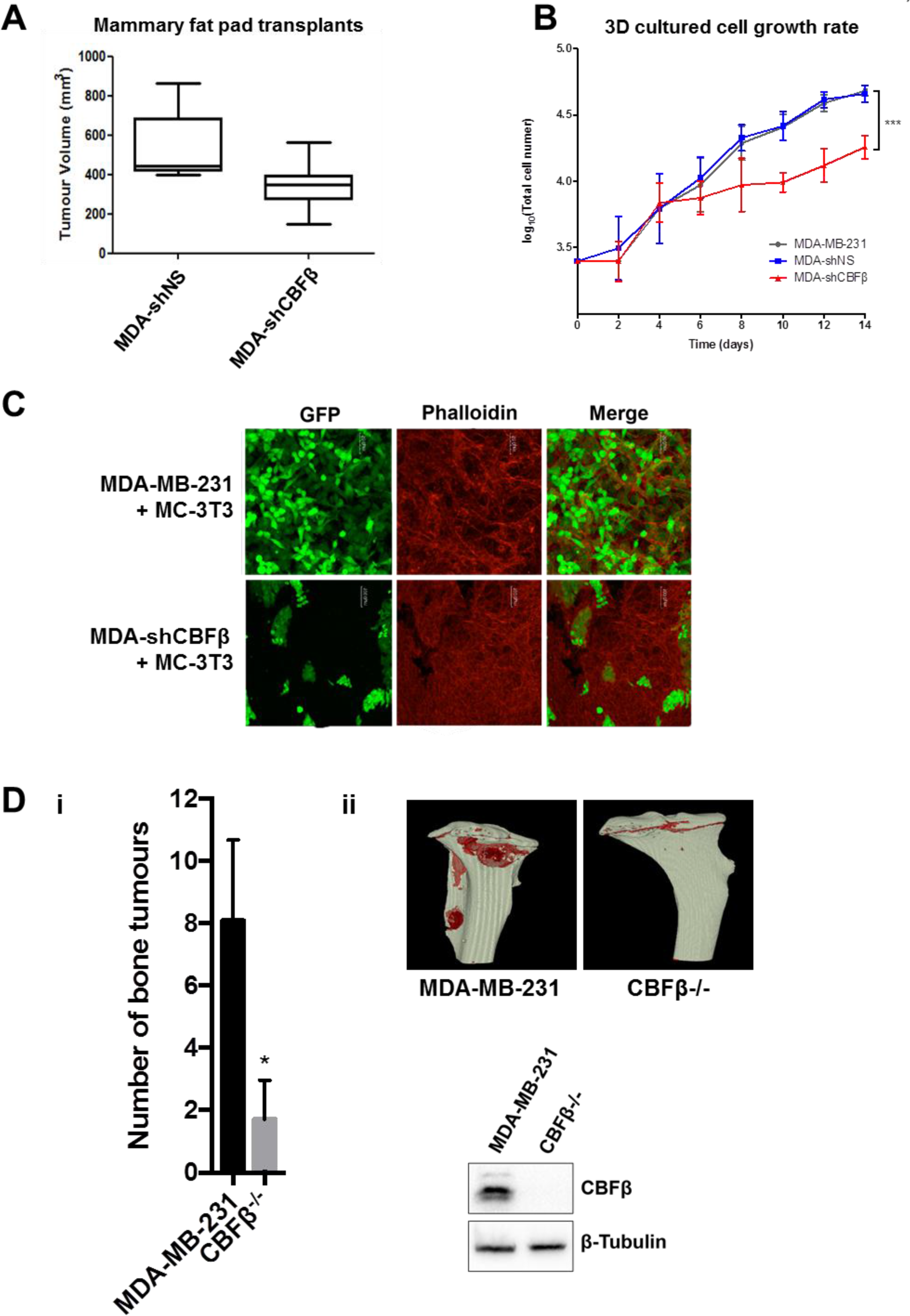
CBFβ contributes to the development of bone metastasis. A) MDA-MB-231 cells were transplanted into the inguinal mammary fat pad of *CD1-Nude* females. Data shown at 4 weeks post-transplantation. Data is presented as mean ± SDM (shNS; n=5; shCBFβ; n=7). The difference between the two groups is significant (p<0.05) as determined using an unpaired Student’s t-test. (B) Growth curves showing knockdown of CBFβ reduces growth rate in 3D culture. MDA-MB-231, MDA-shNS, or MDA-shCBFβ cells were grown on 3D culture and cells were counted every two days. (C) Knockdown of CBFβ inhibits invasion in 3D co-cultures with osteoblasts. 3D cultures of MC-3T3 osteoblasts were grown for 2 months prior to addition of MDAMB-231 or MDA-shCBFβ cells. Confocal images were taken after 8 days of co-culture. Cells were stained with phalloidin. MDA cells were identified by GFP fluorescence. (D) CBFβ silencing reduces tumour growth in bone *in vivo*. MDA-MB-231 control or CBFβ −/− cells were injected i.c. into 6-week old BALB/c nude mice and tumour growth in the hind limbs analysed 26 days later. 3D reconstruction of tumour-bearing tibia showing the presence of osteolytic lesions. The histogram shows the numbers of tumours in bones of each cell line (blue bars: control, red bars: CBFβ knockout).

## Discussion

In this study we have shown that the RUNX co-regulator CBFβ is essential to drive triple-negative breast cancers through EMT. Maintenance of the mesenchymal phenotype is, at least in part, achieved by regulating the expression of the EMT transcription factor Slug. We also demonstrated that the MET induced by loss of CBFβ is completely reversible by re-expression of CBFβ. These findings are important since they demonstrate that, in principle, regulation of RUNX/CBFβ activity can determine the extent to which triple negative breast cancer cells differentiate along the epithelial-mesenchyme continuum. In the context of metastasis *in vivo* this raises the possibility that interactions between cancer cells and the microenvironment influence the activity of RUNX/CBFβ, thereby shifting their phenotype towards the epithelial state and enabling the cells to colonise the new niche. Indeed, the loss of CBFβ significantly reduced the capacity of metastatic cancer cells to invade osteoblast cultures *in vitro* and to form osteolytic lesions *in vivo*.

Previous work has shown that about two thirds of the RUNX1 transcriptome is shared with the RUNX2 transcriptome in MCF7 cells^19^. Our finding that CBFβ, RUNX1 and RUNX2 are all necessary for SNAI2 expression suggests that all three factors combine to ensure a sufficient level of Slug is available to maintain the mesenchymal phenotype. This suggests that none of these factors are redundant in this context. This may reflect the need for a threshold level of RUNX factors to be expressed but does not discount the possibility that RUNX1 and RUNX2 also regulate specific subsets of genes. SNAI2 is a well-established EMT transcription factor that appears to be a key target for RUNX transcription factor complexes and is perhaps one of several genes that contribute to the aggressive nature of triple negative cancers^16 18^.

Mutations in RUNX1 and CBFβ are associated with the ER+ subtype of breast cancer and is therefore predicted to have be a tumour suppressor in this context ^7 8 32^. Indeed, previous studies have shown that RUNX1 suppresses development of ER+ luminal breast cancer but it is not known how CBFβ contributes in this context^33^. In ER-negative MCF10A cells loss of RUNX1 induced EMT, suggesting RUNX1 can act as a pro-metastatic factor^34 20^. This is consistent with our findings that depletion of RUNX1 in MDA-MB-231 cells induces MET. This is also in agreement with the effect of miR-378 expression in MDA-MB-231 cells, in which RUNX1 expression is inhibited and migration and invasion suppressed^35^.

Finally, small molecules inhibitors that inhibit the interaction between CBFβ and RUNX have been shown to inhibit colony formation in a basal-like breast cancer cell line^36^. Our findings that CBFβ is essential to maintain the mesenchyme phenotype, and that it contributes to the formation of bone metastases, suggests that in principle inhibiting this complex might maintain metastatic colonies in a less aggressive epithelial state by driving MET. Thus, targeting the RUNX/ CBFβ complex in this way might be viable option to treat a sub-set of triple-negative breast cancer patients.

## Methods

### Cell lines

Parental MDA-MB-231 expressing GFP were a kind gift from D. Welch, University of Alabama. MDA-MB-231-shCBFb/RUNX1/RUNX2 were produced using Sure Silencing shRNA plasmids (SABiosciences) as previously described^16^. Lines were authenticated by multiplex-PCR assay using the AmpF/STR system (Applied Biosystems) and confirmed as mycoplasma free. Monolayers were grown in complete medium (DMEM/10% FCS/2mmol/L L-glutamine/PenStrep 0.4 μg/mL puromycin, 50 μg/mL geneticin, 500μg/mL hygromycin as required) and maintained in a humidified incubator at 37°C at an atmospheric pressure of 5% (v/v) CO_2_/air.

### Western blotting

Protein was separated on an SDS–PAGE and transferred to Hybond-C Extra nitrocellulose membrane. Primary antibodies included: β-Tubulin (Abcam, ab6046), Lamin-B1 (Abcam, ab16048), CBFβ (Abcam, ab33516), RUNX1 (Abcam, ab23980), RUNX2 (MBL, D130-3), Snai2 (Cell Signalling, C19G7), FLAG (Sigma, F1804). Densitometry was conducted using ImageJ software, which is freely available at http://rsb.info.nih.gov/ij/.

### Cell scratch assay

Confluent monolayers were scratched on day 0 and medium was changed to serum free. Cells were grown in an AS MDW live cell imaging microscope system at 37 °C 5% CO_2_ for 48 hours. Images were taken every 20 minutes and 40 views were taken in each well. Image data analysis was performed using Cell Profile software.

### Overlay three-dimensional culture of breast cells

Matrigel (Corning, 354230) was thawed on ice overnight at 4°C and then spread evenly onto dishes (MatTek P35G-1.0-14-C) or into 24-well plates (Greiner, Bio-one 662892). Cells were resuspended in 3D assay medium (2% Matrigel, 95% DMED, 2%FBS, 1% Pen/Strep, 1% nonessential amino acid, 1% L-glutamine) and plated on to solidified Matrigel. Cells were grown in 5% CO_2_ humidified incubator at 37°C. Assay medium was changed every 3 or 4 days. Cells were fixed at day 14.

### Microscopy

2D culture: Images were collected on a Zeiss Axioimager.D2 upright microscope using a 10x objective and captured using a Coolsnap HQ2 camera (Photometrics) through Micromanager software v1.4.23. Specific band pass filter sets for DAPI, FITC and Cy5 were used to prevent bleed through from one channel to the next.

3D culture: Images were collected on a Leica TCS SP5 AOBS inverted and upright confocal micorscopes. Images were collected using PMT detectors with the following detection mirror settings; [FITC 494-530nm; Texas red 602-665nm; Cy5 640-690nm] using the [488nm (20%), 594nm (100%) and 633nm (100%)] laser lines respectively. When it was not possible to eliminate cross-talk between channels, the images were collected sequentially. When acquiring 3D optical stacks the confocal software was used to determine the optimal number of Z sections. Only the maximum intensity projections of these 3D stacks are shown in the results. Images were then processed and analysed using Fiji ImageJ (http://imagej.net/Fiji/Downloads) [14] which is freely available online.

### Immunofluorescence

For cell grown on coverslips, fixing and permeabilisation was performed in 4 % paraformaldehyde (Sigma) and 0.1 % Triton-100 (Sigma) before blocking in 1 % Bovine Serum Albumin. Primary antibodies included: CBFβ (Abcam, ab33516) and FLAG (Sigma, F1804). Alexafluor secondary antibodies (Invitrogen). The coverslips were mounting on the glass slides using mounting medium with DAPI (Invitrogen P36965). For cells grown in Matrigel, following fixing and permeabilisation as detailed above non-specific staining was blocked using cells were blocked using IF Buffer (7.7mM NaN3, 0.1% bovine serum albumin, 0.2% Triton X-100, 0.05% Tween-20 in PBS) + 10% goat serum. Cells were stained using Phalloidin for F-actin (Sigma, P1951) and mounted using DAPI (Invitrogen).

### Mammosphere culture

Mammosphere culture was carried out as previously describe^37^. Spheres greater than 50µm were counted on day 5.

### Quantitative Reverse Transcription PCR

RNA was extracted using the Qiagen RNAeasy kit according to manufacturer's instructions and quantified on the Nanodrop spectrophotometer (Thermo). Real time one step qRT-PCR was carried out using the QuantiTect SYBR^®^ Green RT-PCR Kit (Qiagen) according to manufacturer’s instructions before analysis on the 7900 PCR machine (Applied Biosystems). A table of the primers used can be found in Supplementary table 1.

### Inducible Cell line production

Mouse CBFβ-FLAG and ER was ligated into pcDNA3.1/Hygro(-) vector producing pcDNA3.1/Hygro(-)-CBFβ-FLAG-ER. Stable lines were made using this vector and cells were transfected using Lipofectamine according to manufacturer’s instructions.

### Nuclear/Cytoplasmic separation

Cells were resuspended in 400μl of ice cold Buffer A (10 mM HEPES pH 7.9, 10 mM KCl, 0.1 mM EDTA, 0.1 mM EGTA, 1 mM DTT, 0.5 mM PMSF) with the addition of complete mini-EDTA-free protease inhibitor cocktail (Roche) and incubated at 4°C for 15 min. Cells were lysed by addition of 10% NP-40 (Sigma) before centrifuging at 4°C and removal of the cytoplasmic extracts in the supernatant. The pellet was then resuspended in ice cold Buffer B (20 mM HEPES pH 7.9, 0.4 M NaCl, 1 mM EDTA, 1 mM EGTA, 1 mM DTT and 1 mM PMSF) containing protease inhibitors and vortexed vigorously for 45 min at 4°C. Nuclear proteins were collected from supernatant following centrifugation at 4°C.

### Invasion Assay

Matrigel Matrix (Corning, 354230) was diluted to final concentration of 300 μg/mL in cold coating buffer (0.01M Tris (pH8.0), 0.7% NaCl) before being added to invasion chambers (Corning Cat, 353097) and left to set overnight at 37°C. 2×10^4^ cells were added to each well in serum free medium and 0.75mL complete medium was to the wells of the wells. Cells were allowed to invade 24 hours in cell culture incubator. Invading cells were fixed and permeabilised with 4% PFA (Electron Microscopy Sciences, 15713-S) and 0.1% Triton (Sigma). Non-Invading cells were removed by a cotton swab. Cells were stained with Crystal violet solution.

### Chromatin immunoprecipitation (ChIP)

ChIP was performed as previously described^38^. ChIP-PCR was performed using Quantitect SYBR green (Qiagen). The primers used can be found in supplementary table 1.

### CRIPSR-Cas9 mediated gene deletion

CBFβ and RUNX1 gene knockout was performed using a double nickase CRISPR-Cas9 strategy as described previously^31^. Guide-RNA sequences were designed using E-CRISP to minimise off target effects^24^. Cells were Fluorescence-activated cell sorted (FACS) for GFP-Cas9 expression 48 hours after transfection and grown up from single colonies prior to genomic DNA PCR and western blot screening.

### Mammary fat pad xenografts

MDA-MB-231 cells were transplanted into the inguinal mammary fat pad of *CD1-Nude* females (Charles River, UK) and tumour volume, measured using callipers, was calculated using the formula ½(length × width^2^). This experiment was carried out in dedicated animal facilities under project licence 60/4181 with adherence to the Animal (Scientific Procedures) Act, the European Directive 2010 and local ethical approval (University of Glasgow).

### Bone tumour growth studies

Tumour growth studies used 6-week-old female BALB/c nude mice (13-16g) (Charles River, Kent, UK). Experiments were carried out in accordance with local guidelines and with Home Office approval under project licence 70/8799, University of Sheffield, UK. Animals were injected with 1×10^5^ MDA-MB-231 control (2014-8-044) or CBFβ CRISPR knockout cells (2015-6-010 CRISPR) via the intra-cardiac (i.c.) route to generate tumours in bone^30^. Animals were culled 26 days following tumour cell injection and hind limbs collected for analyses of tumour growth and associated bone lesions in tibiae and femurs.

### Analysis of bone lesions

Hind limbs were fixed in 4%PFA and scanned by μCT prior to decalcification in 1%PFA/0.5% EDTA and processing for histological sectioning. μCT analysis was carried out using a Skyscan 1272 x-ray-computed μCT scanner (Skyscan, Aartselar, Belgium) equipped with an x-ray tube (voltage, 50kV; current, 200uA) and a 0.5-mm aluminium filter. Pixel size was set to 5.99μm and scanning initiated from the top of the proximal tibia or distal femur. Lytic, tumour-induced bone lesions were counted manually for each bone.

### Statistical analysis

Data is represented as mean +/− SD, n=3 unless otherwise stated.

Statistical significance was measured using parametric testing, assuming equal variance, unless otherwise stated, with standard t-tests for 2 paired samples used to assess difference between test and control samples. * indicates 0.01 < P < 0.05; ** indicates 0.001 < P < 0.01; *** indicates P < 0.001; **** indicates P < 0.0001; N indicates 0.05 < P when compared to control.

## Acknowledgments

The work was funded by Breast Cancer Now (PS, IH and HH), a Cancer Research UK PhD Studentship (PS and HP), a BBSRC PhD studentship and the Manchester Univeristy Alumni Fund (R.A). The Bioimaging Facility microscopes used in this study were purchased with grants from BBSRC, Wellcome and the University of Manchester Strategic Fund. The FACS Aria within the flow cytometry core was purchased with funding from the MRC. Thanks to Michael Jackson for guidance and assistance with sorting.

## Author contributions

RR performed most of the 3D and gene expression analysis of the knockdown cell lines and contributed to the development of the project. HH generated and analysed the CBFβ CRISPR cell line and contributed to the development of the project. RH performed the initial 3D analysis on CBFβ-knockdown cells. NSA generated and analysed the RUNX1 CRISPR cells. HP assisted with CRISPR and writing the manuscript. WD assisted with ChiP assays. AM assisted with the 3D bioreactor cultures. PO, CAF, SMM performed *in vivo* experiments. KB performed in vivo experiments and assisted with writing the manuscript. IH assisted with the conception and design of the project. PS conceived and directed the project, and wrote the manuscript.

## Competing interests

The authors declare that they have no competing interests.

**Materials & Correspondence.** All correspondence should be addressed to PS.

---

**Figure S1.**
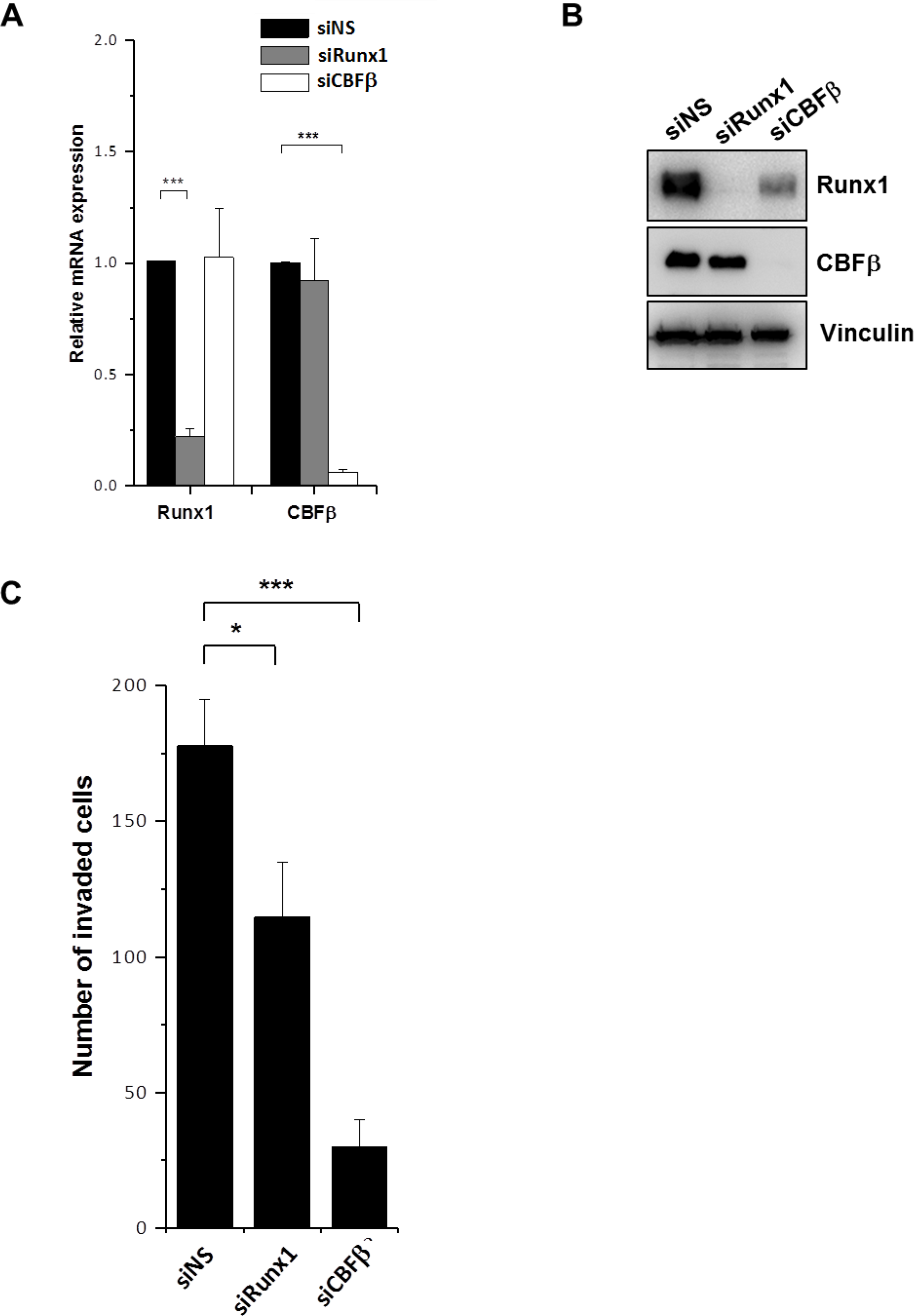
siRNA was used to knockdown RUNX1 and CBFβ in MDA-MB-468 cells, another triple negative breast cancer cell line. 48 hours after transfection, mRNA levels of RUNX1 and CBFβ were quantified (A) and protein (B) to confirm knockdown. Knockdown of CBFβ or RUNX1 leads to a decrease in invasion. Invasive capacity of the cells was assessed *via* Boyden chamber Matrigel assay (C).

**Figure S2.**
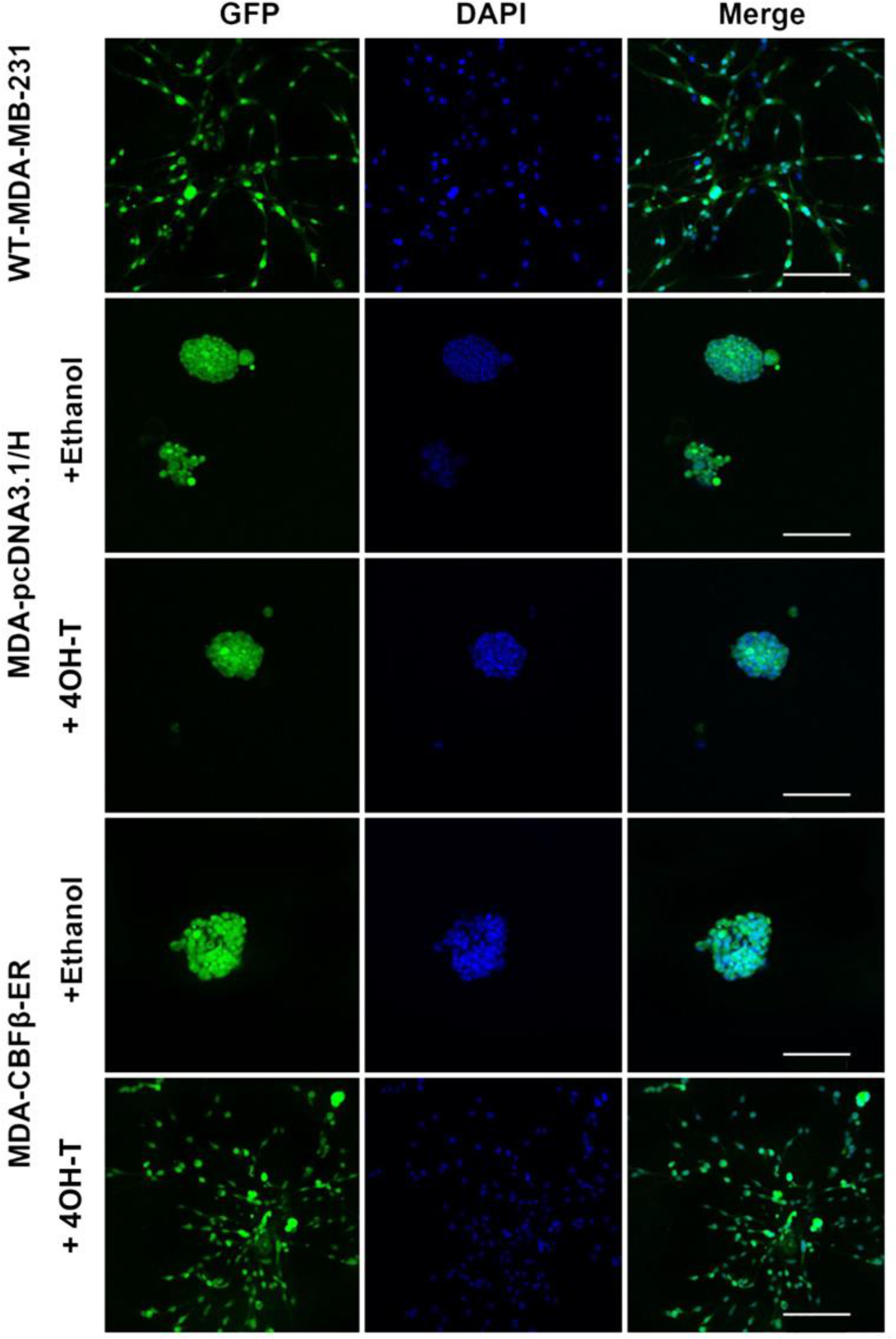
Additional controls for tamoxifen inducible system. Induction of CBFβ-ER with 4OH-T restores the mesenchymal phenotype in 3D culture but no effect is seen in pcDNA3.1/H control cells. The GFP expressing cells were visualised by fluorescence microscopy and DAPI staining. Scale bars are 200μm.

**Figure S3.**
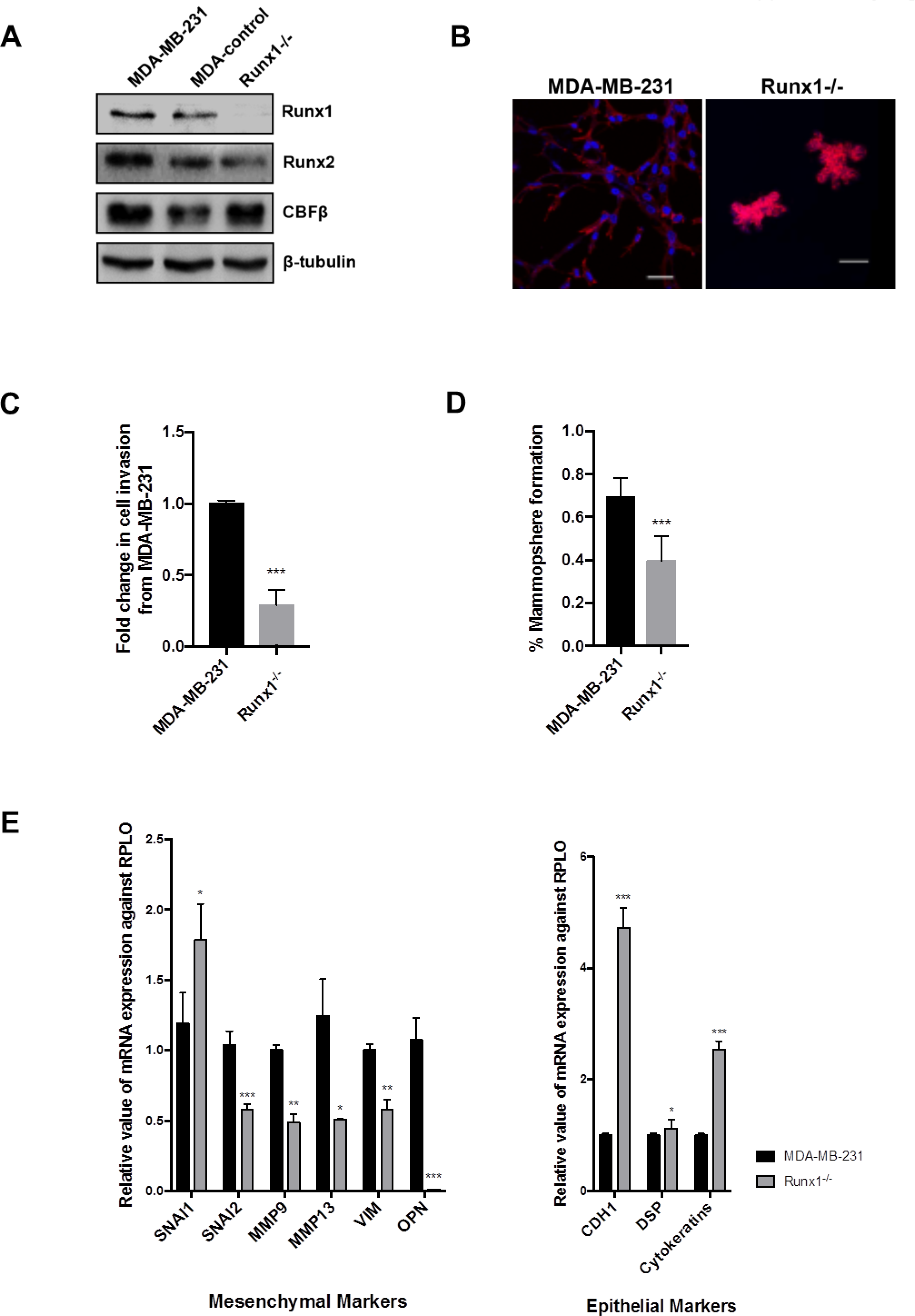
RUNX1 knockout leads to a loss of the mesenchymal phenotype in MDA-MB-231 cells. (A) A double nickase CRISPR-Cas9 system was used to knockout the start of the RUNX1 gene in MDA-MB-231 cells^23^. Two clones with complete RUNX1 loss were generated and a western blot shows the levels of RUNX1, RUNX2 and CBFβ in these cells with β-tubulin used as a loading control. (B) RUNX1 knockout leads to a loss of the mesenchymal phenotype of MDA-MB-231 cells. WT-MDA-MB-231 and RUNX1^−/−^ cells were grown in a 3D Matrigel. Cells were fixed after 14 days and nuclei stained with DAPI (blue) and phalloidin (red). The parental cells showed the stellate mesenchymal pattern after 14 day. RUNX1^−/−^ cells formed discrete clusters. A decrease in invasive capacity of RUNX1 knockout cells was determined using a Boyden chamber Matrigel assay. Scale bars are 50μm (C) A decrease in invasive capacity of RUNX1 knockout cells was determined using a Boyden chamber Matrigel assay. (D) Mammosphere forming efficiency was decreased in RUNX1 knockout cells. Mammospheres were counted after 5 days growth in non-adherent culture. (E) Loss of RUNX1 causes a reduction in mesenchymal markers and increased expression of epithelial markers. RNA from parental MDA-MB-231 and RUNX1^−/−^ cells were subject to qRT-PCR. Acidic ribosomal phosphoprotein P0 (RPLO) mRNA was used for normalisation and relative values of each marker gene mRNA levels are shown.

**Figure S4.**
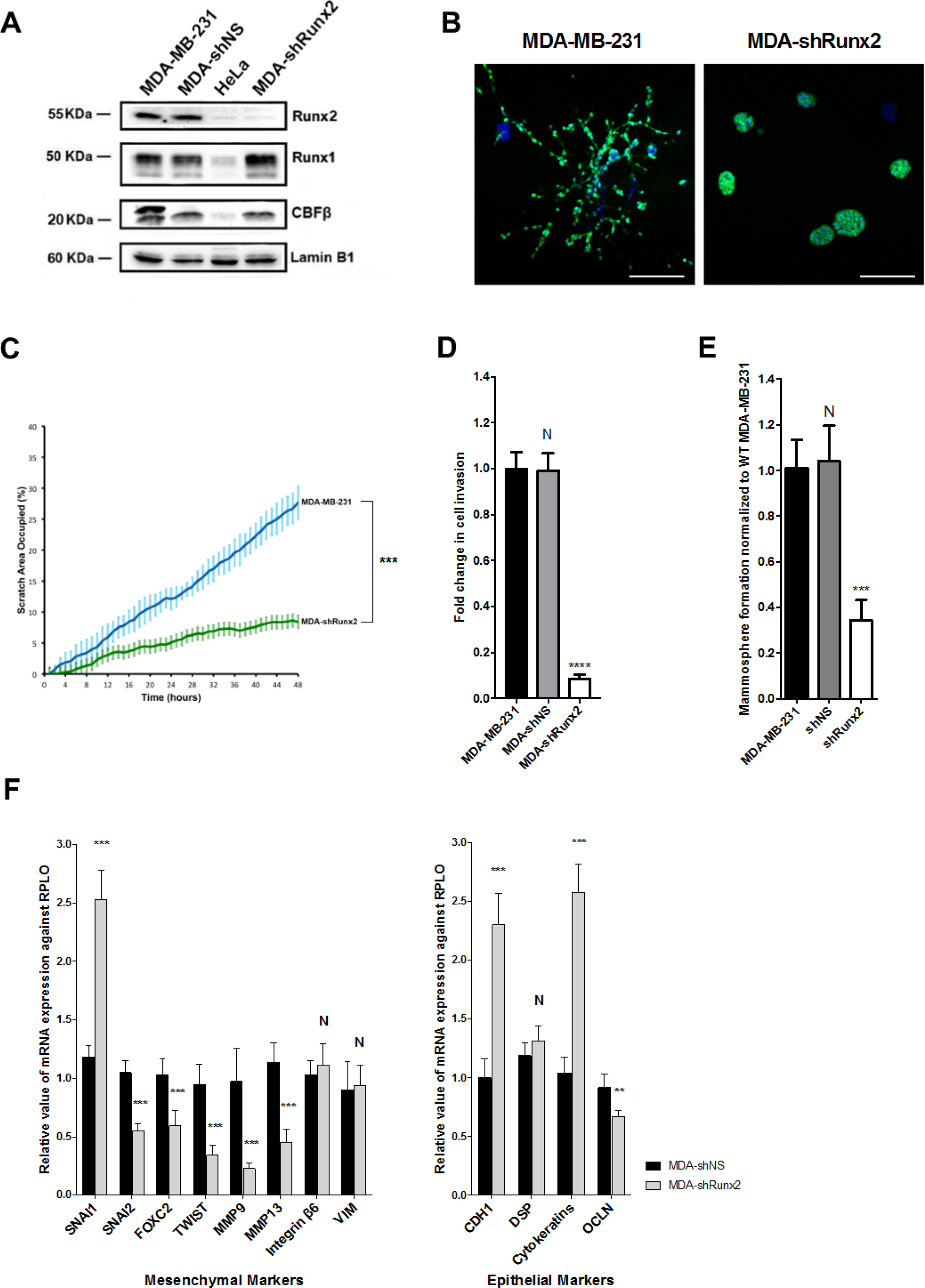
RUNX2 depletion leads to a loss of the mesenchymal phenotype in MDA-MB-231 cells. A) Western blot showing levels of RUNX2, RUNX1 and CBFβ in RUNX2 depleted MDA-MB-231 cells. HeLa cells used as a Runx1/Runx2 low cell line control. Lamin B1 was used as a loading control. (B) WT-MDA-MB-231 and MDA-shRUNX2 cells were grown in 3D Matrigel. Cells were fixed after 14 days and nuclei stained with DAPI (blue). GFP (green) was stably expressed in all cell lines. The parental cells showed the stellate mesenchymal pattern after 14 day. MDA-shRUNX2 cells formed discrete clusters. (C) Scratch assays showing inhibition of migration in MDA-shRUNX2. Live images were taken every 20 mins for 48 hours. (D) Invasive capacity was reduced in shRUNX2 cells. Matrigel invasion assay showing the invasion rates following loss of RUNX2 after 24 hours culture. (E) Mammosphere forming efficiency was also reduced in shRUNX2 cells compared to control cells. Mammospheres were counted after 5 days growth in non-adherent culture. (F) Loss of RUNX2 causes a reduction in mesenchymal markers and increased expression of epithelial markers. qRT-PCR was performed for epithelial and mesenchymal markers in MDA-shNS or MDA-shRUNX2. Acidic ribosomal phosphoprotein P0 (RPLO) mRNA was used for normalisation and relative values of each marker gene mRNA levels are shown.

**Figure S5.**
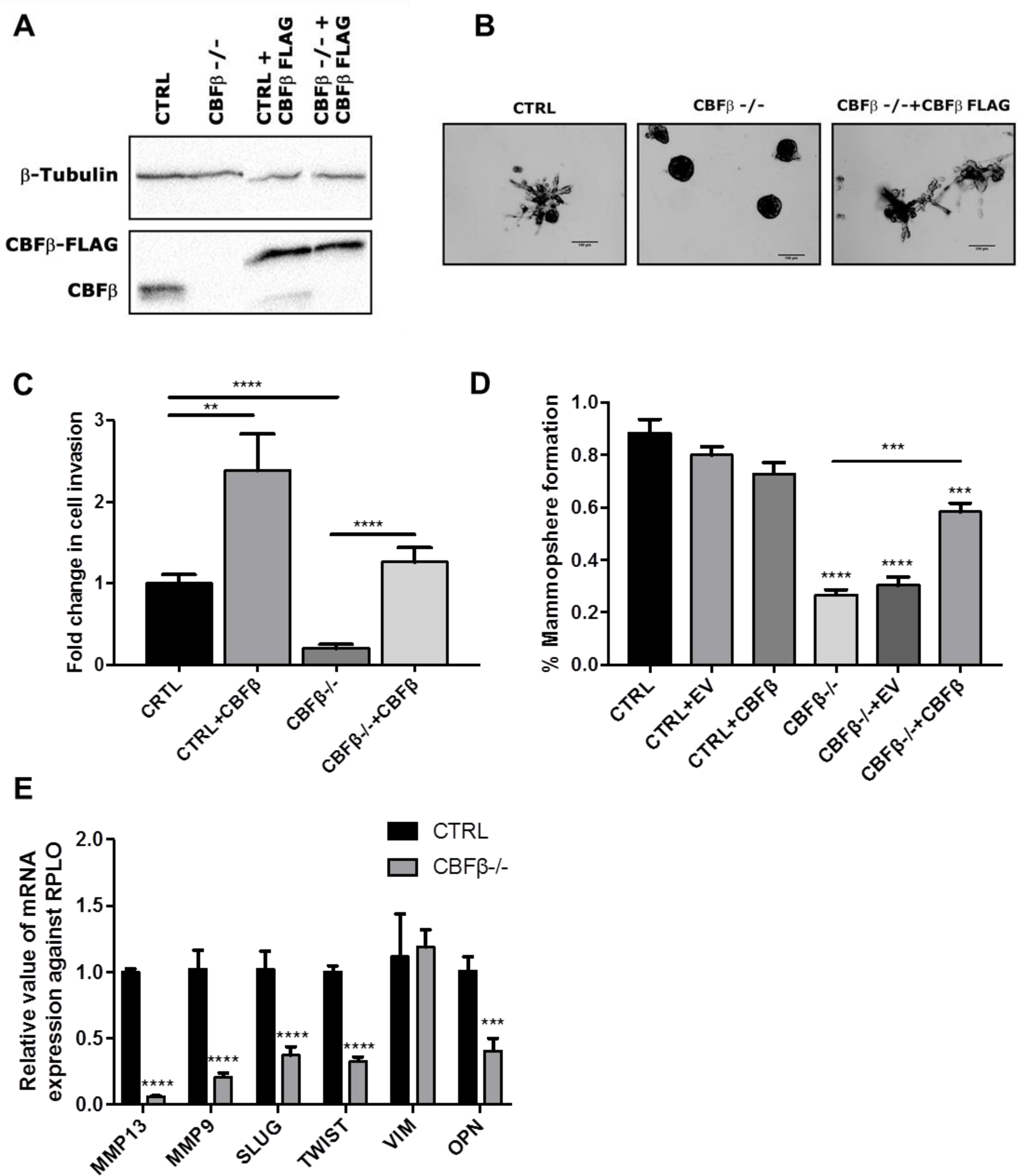
(A) A double nickase CRISPR-Cas9 system (**REF - doi:10.1038/nprot.2013.143**) was used to knockout the start of the CBFβ gene in MDA-MB-231 cells. One clone was generated with complete CBFβ loss and CBFβ-Flag was then re-expressed in this as a rescue. A western blot shows the levels CBFβ in these cells with β-tubulin used as a loading control. (B) CBFβ loss leads to a loss of the mesenchymal phenotype of these cells, which can be rescued *via* CBFβ-Flag re-expression. Cells were grown for 14 days and then imaged using a brightfield microscope. Scale bars are 100μm (C) A decrease in invasive capacity of CBFβ^−/−^ cells was determined using a Boyden chamber Matrigel assay. This can be rescued via CBFβ-Flag re-expression. (D) Mammosphere forming efficiency was also decreased in CBFβ^−/−^ cells and was rescued with re-expression of CBFβ-FLAG. Mammospheres were counted after 5 days growth in non-adherent culture. (E) Loss of CBFβ causes a reduction in mesenchymal markers. RNA from MDA-MB-231 control and CBFβ^−/−^ cells were subject to qRT-PCR. Acidic ribosomal phosphoprotein P0 (RPLO) mRNA was used for normalisation and relative values of each marker gene mRNA levels are shown.

**Supplementary Table 1.**
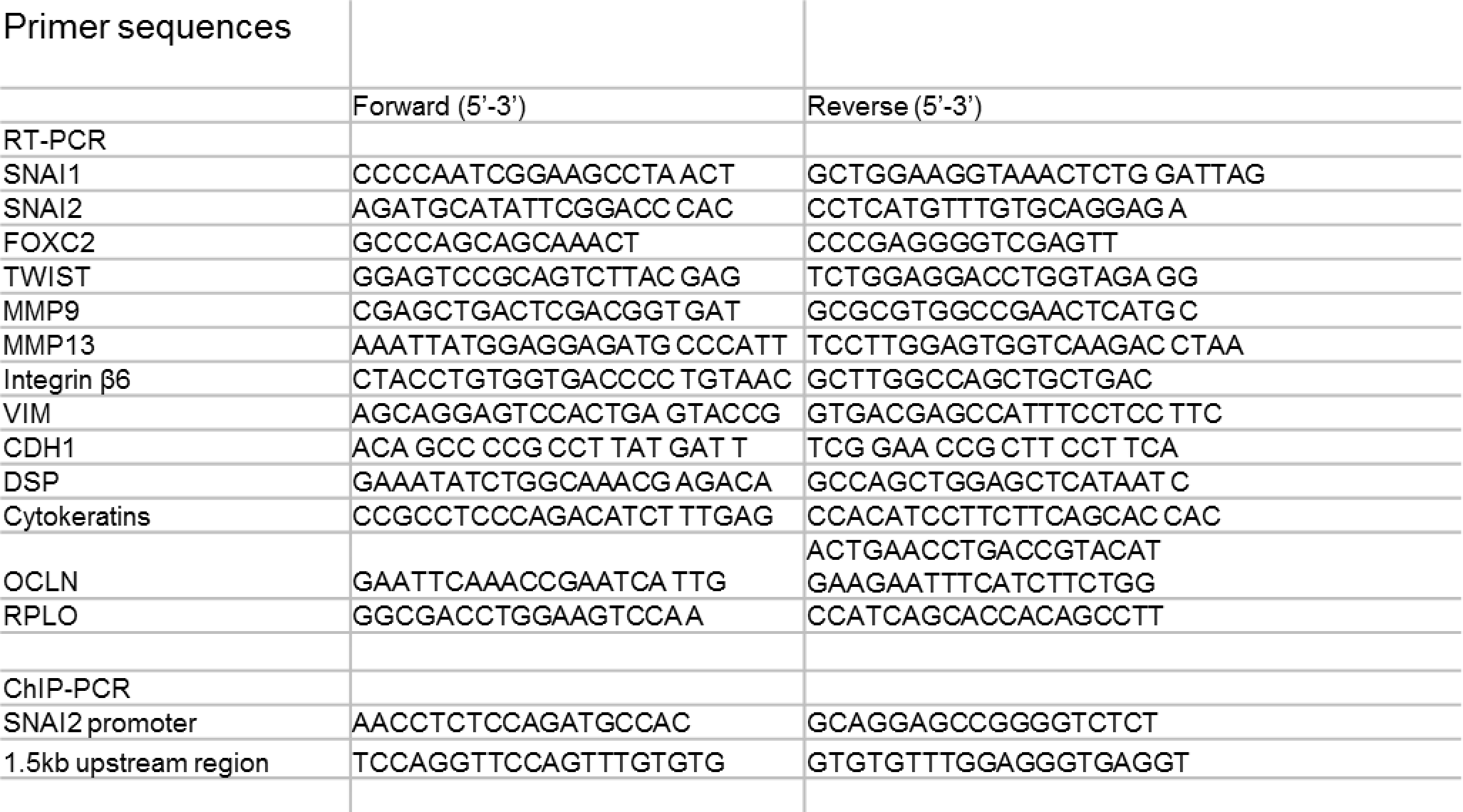

